# Camphor white oil induces tumor regression through cytotoxic T cell-dependent mechanisms

**DOI:** 10.1101/386789

**Authors:** Yalda Moayedi, Sophie A Greenberg, Blair A Jenkins, Kara L Marshall, Lina V Dimitrov, Aislyn M Nelson, David M Owens, Ellen A Lumpkin

## Abstract

Bioactive derivatives from the camphor laurel tree, *Cinnamomum camphora*, are posited to exhibit chemopreventive properties but the efficacy and mechanism of these natural products have not been established. We tested an essential-oil derivative, camphor white oil (CWO), for anti-tumor activity in a mouse model of keratinocyte-derived skin cancer. Daily topical treatment with CWO induced dramatic regression of pre-malignant skin tumors and a two-fold reduction in cutaneous squamous cell carcinomas. We next investigated underlying cellular and molecular mechanisms. In cultured keratinocytes, CWO stimulated calcium signaling, resulting in calcineurin-dependent activation of nuclear factor of activated T cells (NFAT). *In vivo*, CWO induced transcriptional changes in immune-related genes, resulting in cytotoxic T cell-dependent tumor regression. Finally, we identified chemical constituents of CWO that recapitulated effects of the admixture. Together, these studies identify T cell-mediated tumor regression as the mechanism through which a plant-derived essential oil diminishes established tumor burden.

**SUMMARY BLURB:** Essential oil derived from the camphor tree acts by stimulating immune cell-dependent regression of skin tumors in a mouse model of cutaneous squamous cell carcinoma.

## Introduction

Plant-derived essential oils have been used since the Middle Ages for their medicinal and antiseptic properties (Bakkali et al., 2008). In recent years, essential oils have garnered heightened interest for their therapeutic value in treating human ailments; however, the efficacy of such treatments and mechanisms of action have rarely been tested in controlled studies. Camphor white oil (CWO), which is produced by steam distillation of wood from the camphor laurel tree (*Cinnamomum camphora*), is marketed as a fragrance and herbal remedy with antifungal, antiseptic and medicinal properties including circulatory stimulation, increased metabolism and improved digestion (Lee et al., 2006, Satyal et al., 2013, Yang et al., 2014). CWO primarily consist of a mixture of structurally related terpenes, a class of chemical compounds composed of isoprene units with distinct aromatic qualities that plants generate for defensive purposes.

CWO’s terpene constituents are widely found in natural and industrial products. These terpenes are volatile, bioactive and readily absorbed by the skin. Abundant constituents of CWO include eucalyptol, a natural cough suppressant, and limonene, an antiseptic. Counterintuitively, camphor is found at only a trace amounts in this distillate. Interestingly, several of the purified components of CWO, or plants that contain these ingredients, are proposed to have anti-tumorigenic effects; however, few of these compounds has been rigorously tested *in vivo* for these properties (Bhattacharjee & Chatterjee, 2013, Chaudhary et al., 2012, Chidambara Murthy et al., 2012, Kusuhara et al., 2012, Lee, Hyun, 2006, Russin et al., 1989). Terpene constituents of CWO are also common in skin-care products; however, mechanisms through which these bioactive compounds act on epithelial cells have not been identified.

Cutaneous squamous cell carcinoma (cSCC), a non-melanoma skin cancer, is the most common form of skin cancer with metastatic potential (Bachelor et al., 2011). cSCC is often preceded by pre-malignant skin lesions termed actinic keratoses (AKs) (Dodds et al., 2014). AKs are generally treated by field-directed therapy; however, they can recur and convert to cSCC with metastatic potential, particularly in organ transplant recipients and patients with other risk factors such as smoking (Leonardi-Bee et al., 2012, Yu et al., 2014). Thus, there is a need for novel, non-toxic and cost-effective therapeutics to prevent malignant conversions in high-risk patients. Remarkably, nearly 50% of all drugs approved for use in cancer since the 1940s are either natural products or derivatives, suggesting that essential oils such as CWO could be a source for developing new therapeutics (Newman & Cragg, 2016).

To identify whether CWO has biological effects on keratinocyte-derived skin lesions, we employed a pre-clinical model of cSCC in mice. Our results show that daily topical application of naturally derived CWO induces premalignant tumor regression and reduces malignant conversion to cSCC *in vivo*. CWO induced calcium/calcineurin mediated Nuclear Factor of Activated T cells (NFAT) translocation in keratinocytes, causing transcriptional effects that culminate in immune system activation and cytotoxic T cell-dependent tumor regression. Finally, two purified constituents, α-pinene and d,l-limonene, recapitulated the anti-tumor effects of the essential oil. Together, these studies identify CWO as potent anti-tumor agent, establish underlying biological mechanisms and pinpoint lead compounds for further development as new treatments for cSCC.

## Results

### CWO induces cutaneous tumor regressions

To test whether CWO possesses anti-tumor activity *in vivo*, a two-step chemical carcinogenesis model of cSCC was employed in mice (Abel et al., 2009). Tumor formation was initiated by a single topical application of 7,12-Dimethylbenz[a]anthracene (DMBA) to the dorsal skin of mice, followed by twice weekly applications of the tumor promotor 12-O-tetradecanoylphorbol-13-acetate (TPA) for 12 weeks (**Fig. 1A**). In this model, premalignant lesions called papillomas form, up to 50% of which will convert to cSCCs, depending on mouse strain and dosage of initiator and promotor (Abel, Angel, 2009, Hennings et al., 1993, Owens et al., 1999). Chemically induced tumors in mice have both histological and molecular profiles that recapitulate SCCs in humans (Abel, Angel, 2009, Nassar et al., 2015).

**Fig. 1.**
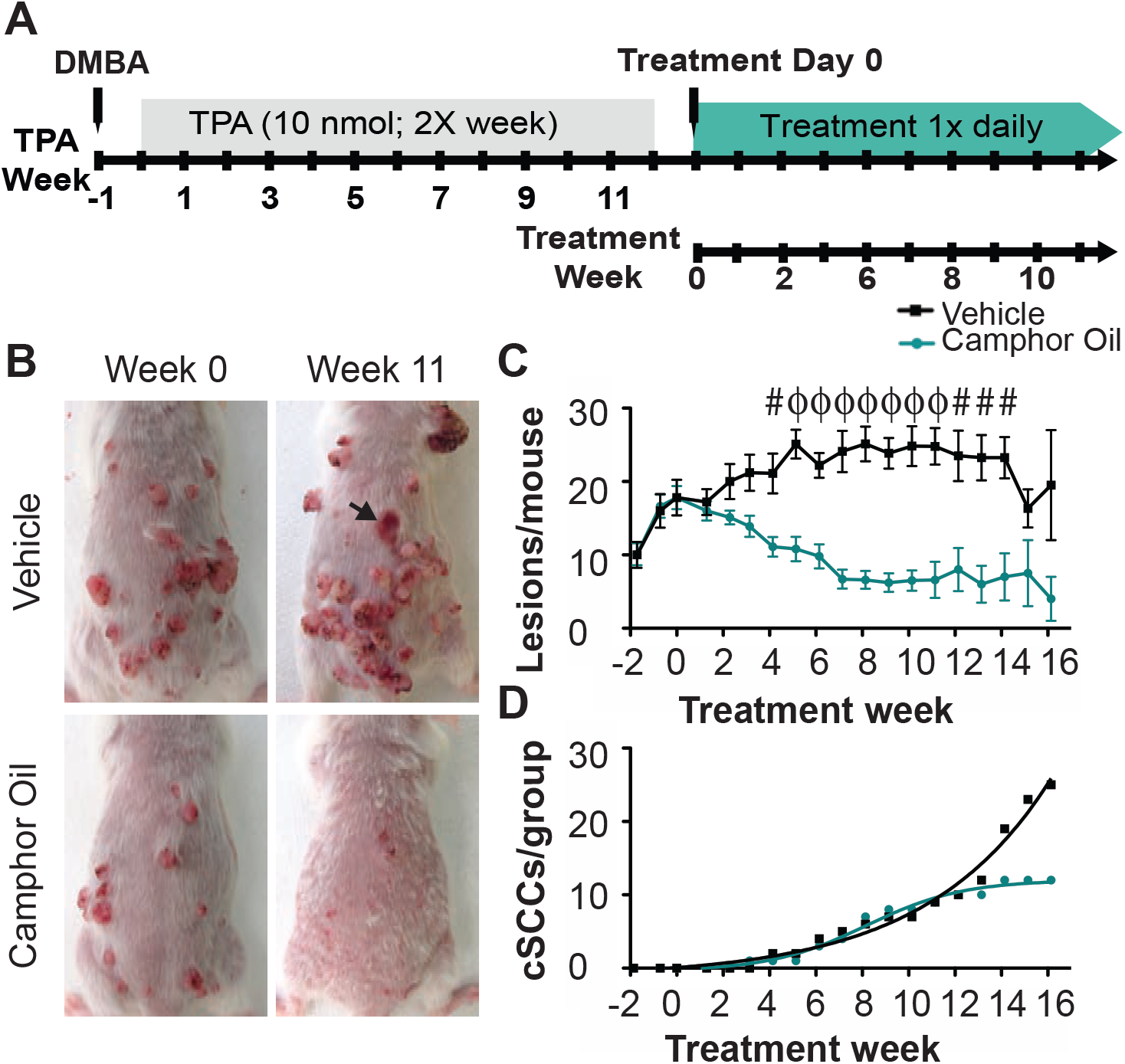
CWO treatment reduces skin tumor burden and decreases malignant conversion to cSCC. **A.** Schematic of the two-step chemical carcinogenesis model. **B.** The same mice are shown at week 0 and week 11. Tumors progressed in size and severity in the vehicle treated mouse (wk 0=16 vs wk 11=28 tumors). CWO treated tumors regressed (wk 0=12 tumors vs wk 11=0 tumors). Arrow denotes a malignant conversion in the vehicle treated group. **C.** Topical CWO treatment significantly reduced the average number of lesions per mouse. Twoway ANOVA [W=10 mice/group, P<0.0001, group effect F(1,257)=196.04]. ^#^P <0.01, ^ϕ^P <0.0001 Bonferroni post hoc. **D.** Malignant conversion to SCC showed significant difference between treatment groups (Boltzmann fits, difference between fits F(4,30)=46.28, P<0.0001; CWO: R^2^=0.99, Max cSCC=12.0 weeks; Vehicle: R^2^=0.98, Max csCc=15046). See **Fig1-S1 & 1-S2** for information on a second cohort. Histological analysis of tumors are in **Fig1-S3**.

After papillomas developed, mice were randomly assigned to one of two treatment groups (N=10 mice per group; CWO treated: 18.2±5.4 lesions per mouse [mean±SD]; vehicle control: 18.7±7.9 lesions per mouse; P=0.87, Student's two-tailed *t* test). Mice then received daily topical treatments of 20% CWO in acetone vehicle or vehicle alone for up to 24 weeks. Vehicle-treated mice showed a stable lesion burden up to 13 weeks after treatment onset, where existing lesions increased in size and tumor grade over time (**Fig. 1B-C**). At 14 weeks, lesion burden (**Fig. 1C**) decreased in the control group because mice with the highest tumor burden reached experimental endpoints. By contrast, daily topical CWO induced dramatic regression of lesions in size and number, which was apparent within two weeks (**Fig. 1B-C**). Remarkably, camphor-oil treatment resulted in a nearly two-fold decrease in the incidence of malignant cSCCs by 16 weeks (**Fig. 1D**). Although the number of malignant conversions reached maximum at twelve weeks of camphor-oil treatment, conversions in the vehicle treated group continued to rise until all animals in that group reached experimental endpoint (16 weeks, see Extended Experimental Methods for endpoint criteria). Similar results were found in a second independent cohort (**Fig. 1-S1 and S2**). Although median survival curves were comparable between treatment groups, a subset of camphor-oil treated individuals showed a 39% increase in survival times compared with vehicle (**Fig. 1-S1D-E**). Tumors from vehicle and CWO-treated mice were histologically similar (**Fig. 1-S3**), whereas areas where lesions regressed in CWO treated mice resembled hyperproliferative skin (**Fig. 1-S3I-J**). Thus, we conclude that CWO has robust anti-tumor activity on keratinocyte-derived lesions *in vivo*.

### CWO induces calcium/calmodulin-dependent NFAT signaling in keratinocytes

To define mechanisms that might underlie this striking reduction in tumor burden, we first sought to uncover molecular pathways through which CWO exerts bioactive effects on keratinocytes. We reasoned that CWO might activate Nuclear Factor of Activated T cells (NFAT) in keratinocytes. In a low calcium environment, this transcription factor localizes to the cytoplasm. In response to cytoplasmic calcium signaling, NFAT is dephosphorylated by calcineurin, a calcium/calmodulin-dependent phosphatase, which allows NFAT to enter the nucleus and direct target gene expression (Crabtree & Olson, 2002, Hannanta-Anan & Chow, 2016, Li et al., 1998). Three lines of evidence focused our attention on NFAT signaling. First, previous studies have shown that terpenes activate calcium signaling in mammalian cells *in vitro* (Oz et al., 2015, Rodrigues et al., 2016, Vogt-Eisele et al., 2007), which is an essential step in NFAT activation. Second, inhibition of calcineurin with cyclosporine A (CSA) promotes cSCC in humans and animal models, underscoring the role of calcineurin/NFAT in SCC pathogenesis (Euvrard et al., 2003). Finally, CSA regulates the cell cycle in SCC keratinocytes *in vitro*, suggesting that calcineurin/NFAT signaling has direct effects on keratinocyte relevant to cancer biology (Dotto, 2011, Wu et al., 2010).

We first asked whether CWO induces calcium signaling. Normal human epidermal keratinocytes were loaded with the ratiometric calcium indicator Fura-2 AM and responses to CWO application were monitored with live-cell imaging. CWO induced rapid increases in cytoplasmic calcium in a dose-dependent manner (**Fig. 2A-C**). In many cases, calcium waves were observed in individual keratinocytes (**Fig. 2-S1**), an activation pattern that induces the calcium/calcineurin-dependent transcription factor NFAT to translocate to the nucleus and direct expression of target genes (Crabtree & Olson, 2002, Hannanta-Anan & Chow, 2016, Li, Llopis, 1998).

**Fig. 2.**
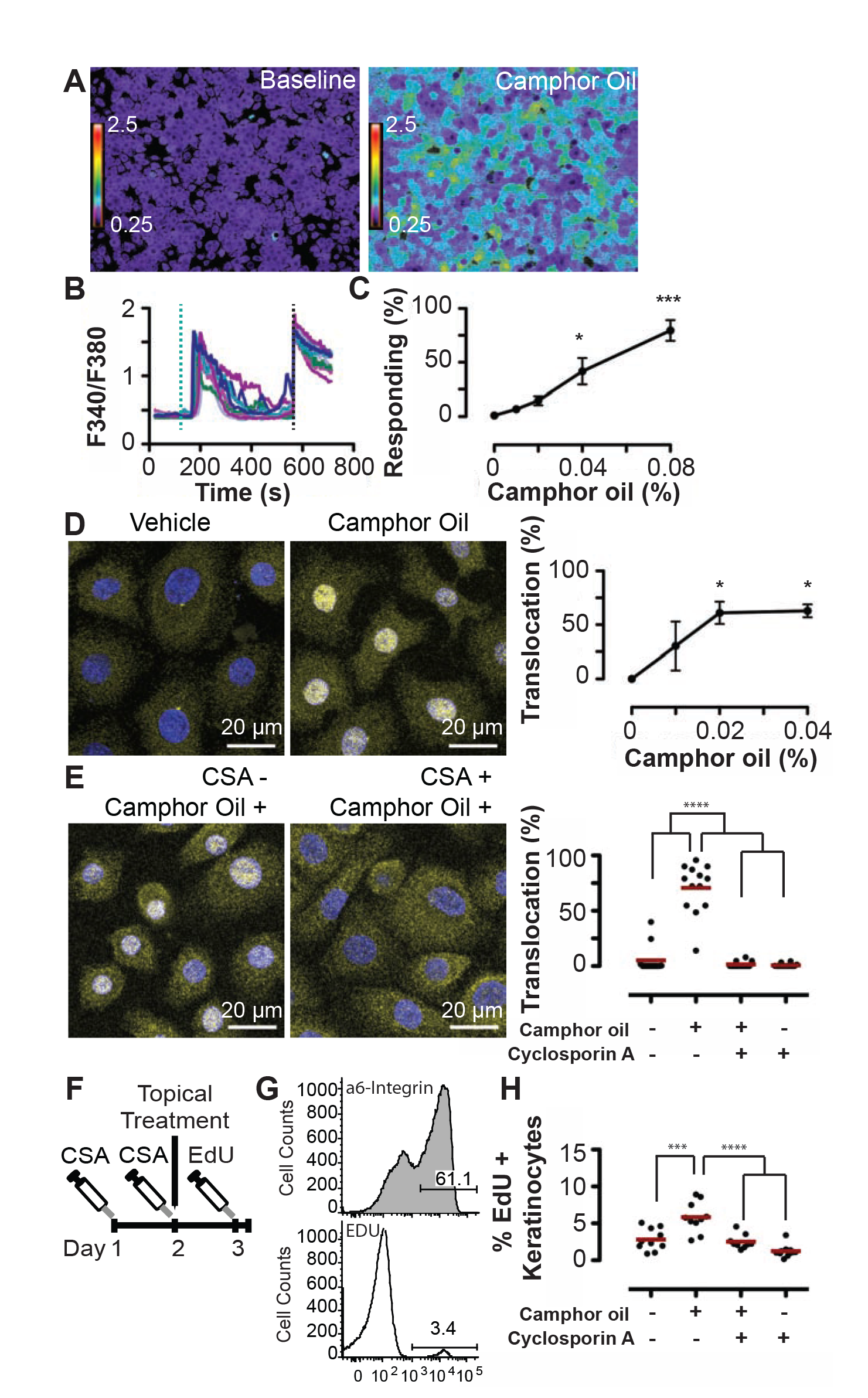
CWO induces keratinocyte proliferation dependent on calcium/calcineurin. CWO induced calcium signaling in human epidermal keratinocytes in vitro (**A-C**). **A.** Keratinocytes before and after 0.04% CWO application. **B.** Calcium transients with 0.04% CWO application. Teal dashed line indicates CWO application, black dashed line indicates positive control (histamine). See **Fig 2-S1** for examples of individual traces. **C.** Percent of cells responding to CWO: 0.04–0.08% CWO significantly induced calcium responses (N=7–10 experiments/group from four individual keratinocyte donors P<0.0001, One-way ANOVA, Bonferroni’s post-hoc comparison with DMSO vehicle (Veh). See **Fig 2-S2** for viability analysis. **D.** Cytosolic and nuclear localization of NFAT (yellow) and DAPI (blue) with 0% and 0.02% CwO. (**Right**) Percent of keratinocytes with nuclear NFAT localization following incubation with 0–0.04% CWO (n=3 experiments with >1200 cells per replicate; P<0.05 One-way ANOVA, Bonferroni’s post-hoc comparison with Veh). **E.** Cells were pre-treated for 1 h with CSA or vehicle followed by a 30-minute treatment with 0.04% CWO or vehicle. NFAT localization after incubation with Veh/Veh, Veh/0.04% CWO (**Left**), 1 μM CSA/0.04% CWO, and 1 μM CSA/Veh (**Right**). Two-way ANOVA found a significant effect of CWO (P<0.0001) and CSA (P<0.001). Bonferroni post-hoc comparison revealed a significant difference between Veh/Veh and Veh/CWO samples that was abolished with CSA pre-treatment. N=10–11 experiments/group. One statistical outlier (as determined by Grubb’s test) was removed from both CSA/V and CSA/CWO groups. **F.** Mice were treated 2x with CSA or Veh every 24 h followed by one application of CWO or Veh. 23 h later, mice were given a single injection of EdU. After a 1 h, keratinocytes were collected. **G.** Representative flow gates. The percentage of EdU+ basal keratinocytes (α6-integrin+) were quantified. **H.** CWO induced a significant increase in basal keratinocyte proliferation. Two-way ANOVA P_CSA_<0.001, P_CWO_<0.0001. Bonferonni post hoc analysis revealed a significant difference between Veh/CWO and Veh/Veh treatment groups, which was abolished by CSA pre-treatment. N=9–10 animals/group, one statistical outlier was removed from the CSA/CO group. * P<0.05, *** P<0.001, ****P<0.0001, Red lines indicate group mean values.

To test the hypothesis that NFAT is activated downstream of CWO treatment in keratinocytes, we analyzed nuclear translocation of NFATc1, an isoform with well documented activity in human keratinocytes (**Fig. 2D**)(Horsley et al., 2008, Mammucari et al., 2005, Santini et al., 2001, Tripathi et al., 2014). Keratinocytes treated *in vitro* with 0.02–0.04% CWO displayed significant translocation of NFATc1 without compromising cell viability (**Fig. 2-S2**). This effect was completely eliminated by pre-treatment with CSA, indicating that NFAT translocation was dependent on calcium/calcineurin signaling (**Fig. 2E**). Thus, we conclude that CWO induces NFAT translocation through calcium/calcineurin signaling in keratinocytes.

NFAT activity has context-dependent effects on keratinocyte proliferation, both maintaining stem cell quiescence and inducing keratinocyte proliferation (Goldstein et al., 2014, Horsley, Aliprantis, 2008, Keyes et al., 2013, Tripathi, Wang, 2014). We postulated that CWO might also alter the cell cycle through NFAT activity. We noted that CWO application induces mild thickening of the mouse epidermis adjacent to treated tumors, which suggests that CWO induces proliferation. To quantify proliferation and its dependence on calcium/calcineurin/NFAT signaling, we treated normal mice with a single topical application of CWO or vehicle in conjunction with either CSA or vehicle treatment (**Fig. 2F**). Twenty-four hours after treatment, a single 1-h pulse of EdU was administered and keratinocytes were harvested. The fraction of EdU+ keratinocytes in the proliferative basal layer was measured with flow cytometry (Blanpain & Fuchs, 2006). Keratinocyte proliferation was enhanced two-fold by CWO treatment and this effect was blocked by CSA treatment (**Fig. 2G–H**). We conclude that CWO induces calcium/calcineurin/NFAT signaling that mediates biological effects on keratinocytes *in vivo*.

Given that CWO induces calcium signaling and proliferation in keratinocytes, we hypothesized that it might work through TRPV3, a calcium channel which is expressed in keratinocytes and is activated by camphor and related terpenes (Borbiro et al., 2011, Vogt-Eisele, Weber, 2007). To test whether TRPV3 is required for CWO’s effects in the skin, we performed *in vivo* proliferation assays on TRPV3 knockout animals and wild-type littermates (**Fig. 3**). TRPV3 disruption had no effect on CWO-meditated proliferation; therefore, we conclude that the effects of CWO are not TRPV3 dependent.

**Fig. 3.**
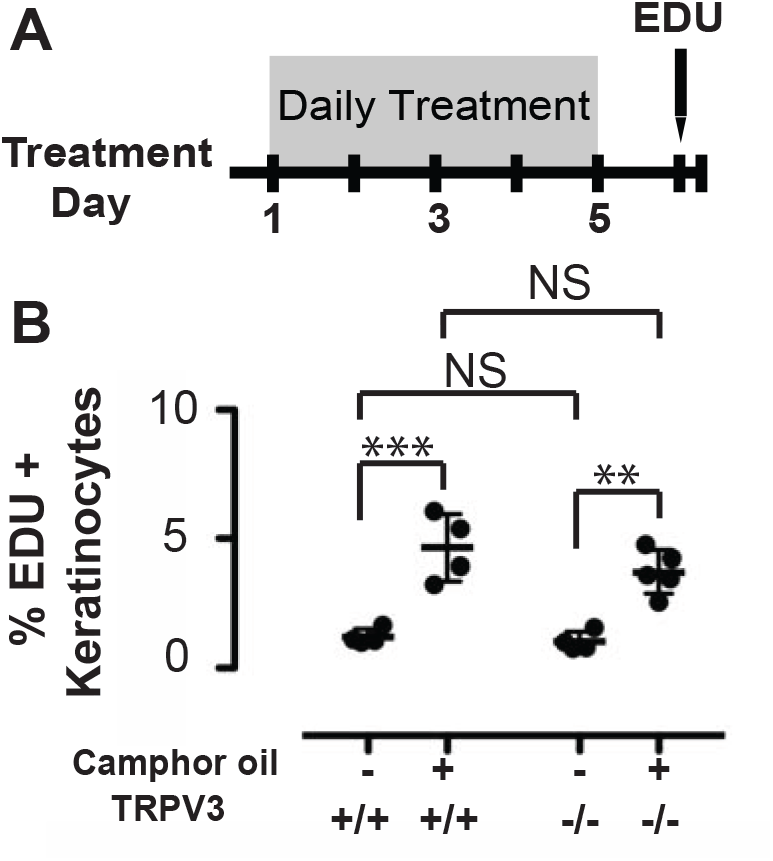
Camphor-oil’s biological effects on keratinocytes is independent of TrpV3. **A.** TrpV3 mutant and age-matched wild-type mice were treated with CWO or vehicle control for 5 days. Mice were then given a 1-h pulse of EdU and keratinocytes were collected and analyzed as in Fig. 2. **B.** CWO induced a significant increase in basal keratinocyte proliferation; however, there was no significant effect of TRPV3 knockout on proliferation (N=4–5 mice per group; Two-way ANOVA, non significant genotype effect, treatment effect: p<0.0001; Bonferroni post hoc analysis: **p<0.01,***p<0.001).

### CWO treatment induces immune-dependent tumor clearance

We next took an unbiased, genome-wide approach to identify genetic pathways involved in CWO-induced tumor regression. To this end, we performed RNA-sequencing of CWO-treated mouse tissues. Samples were harvested from pre-malignant tumors collected after six weeks of treatment with vehicle or CWO (N=3 mice per group, 4 pooled tumors per mouse). To identify targets directly downstream of CWO activity in the skin, RNA-sequencing was also performed on epidermis isolated 24 h after a single treatment with CWO or vehicle. We identified 293 genes differentially expressed in CWO-treated tumors and 1677 transcripts altered in CWO-treated epidermis (**Fig. 4A**). Seventy-seven genes were common in both datasets. We compared biological processes significantly altered in both tissues by CWO treatment. These included several processes that influence tumor biology, including blood vessel/vasculature development, skin development and cell death (**Fig. 4A**). Interestingly, several enriched nodes were consistent with an effect on immune regulation including: cellular response to cytokines, granulocyte migration, and leukocyte chemotaxis. Based on this intriguing finding, we postulated that CWO treatment of keratinocytes stimulates immune-cell recruitment, a mechanism that could lead to engulfment of tumor cells. To explore this possibility, we analyzed gene ontology (GO) terms enriched in each tissue group separately with a focus on immune pathways (**Fig. 4-S1**). CWO treatment of tumors stimulated inflammatory response and chemotaxis pathways. Additionally, myeloid leukocyte migration and T cell migration pathways were significantly enriched in epidermis. These findings, along with the striking effect on regression of existing tumors, suggest that CWO stimulates a T cell-mediated immune responses.

**Fig. 4.**
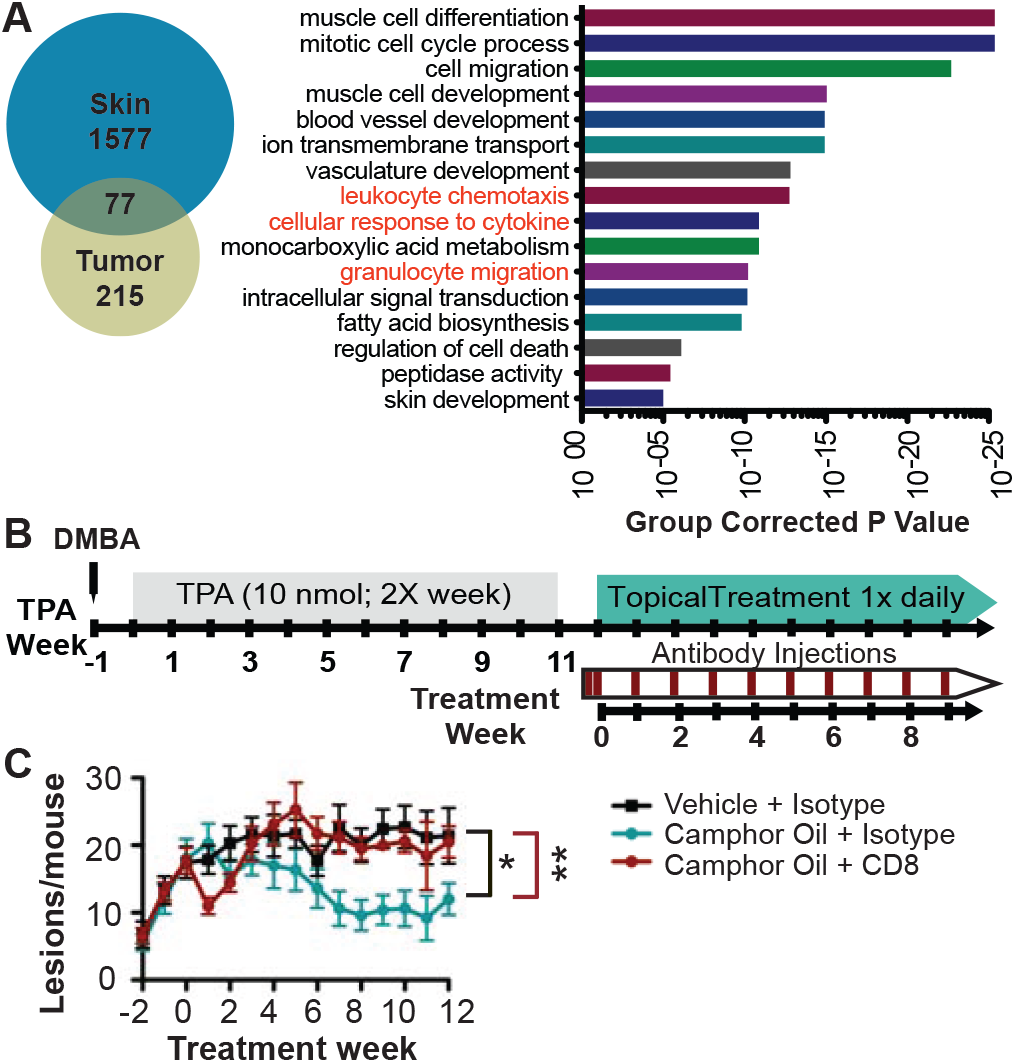
CWO-induced tumor regression requires cytotoxic T cells. **A.** Epidermis and tumors treated with CWO or vehicle were analyzed with RNA sequencing and results were compared (left). Epidermis was treated with a single application of CWO, tumors were collected after six wks of treatment. 1577 genes were differentially enriched in skin, 215 in tumors, and 77 in both groups with CWO treatment. GO-analysis of terms enriched with CWO treatment in both papillomas and keratinocytes were analyzed (right) and compared to identify common pathways affected in both groups. Terms consistent with immune activation are in red font. See **Fig. 4-S1** for GO terms from individual groups. **B.** Tumors were induced as before, mice were split into matched cohorts, then injected with T-cell blocking or isotype control antibodies on day -1, 2 and then then 1x weekly. **C.** CD8 blocking antibody reversed the effects of CWO treatment (*N*=7 mice/group, two-way ANOVA identified a significant interaction between time and treatment group (P<0.05), and significant effects of both time and treatment group (P<0.0001). One-way ANOVA split by treatment group, P<0.01, Tukey’s multiple comparison test *P<0.05, **<0.01. See **Fig. 4-S2** for information on antibody blocking and CD4 experiments.

To dissect the dependence of CWO’s anti-tumor effects on immune substrates, we tested the necessity for CD4+ and CD8+ T cells, which are lymphocyte subsets commonly associated with anti-tumor immunity. CD4+ T helper (T_H_) cells activate and promote immune responses by releasing cytokines, priming CD8+ T cells, and activating antigen presenting cells. Upon activation, CD8+ cytotoxic T cells directly kill cancerous cells (Shiku, 2003). Tumors were chemically induced and then mice were randomly assigned to one of four treatment groups: vehicle with isotype control injections, CWO with isotype control injections, CWO with CD8 blocking antibody, and CWO with CD4 blocking antibody. Mice received daily topical treatments with CWO or vehicle along with weekly injections with antibodies or isotype (IgG) controls (**Fig. 4B**). The effects of CWO treatment were completely reversed by administering a CD8-blocking monoclonal antibody (**Fig. 4C**), which was effective in reducing CD8+ circulating cells up to 6 days after each application (**Fig. 4-S2A**). CD4 blocking antibodies appeared to have an intermediate effect on CWO activity (**Fig. 4-S2B**); however, CD4+ cells were present at sacrificing, indicating that this population was not fully depleted by antibody treatment (**Fig. 4-S2**). Collectively, these data suggest that CWO-mediated tumor regression is dependent on CD8+ cytotoxic T cells.

### Identification of CWO’s active ingredients and effective concentrations

Finally, we sought to determine active compounds in CWO that stimulate tumor regression. As a natural plant derivative, CWO’s mixture of terpenes varies depending on terroir (Satyal, Paudel, 2013). Using gas chromatography-mass spec (GC-MS), we found that many structurally related terpenes were present in CWO, with eucalyptol and d,l-limonene being the most abundant constituents (**Fig. 5A**). As expected due to distillation, camphor was identified only in trace quantities in CWO and is thus not expected to be an active ingredient. To test whether individual constituents recapitulate the anti-tumor effects of CWO, tumors were induced as before and matched cohorts were treated with one of five terpenes (eucalyptol, d,l-limonene, α-pinene, g-terpinene, and camphene; N=5 mice per group). Compounds were chosen based on abundance in CWO, confidence of identification in GC-MS (>90% matching to reference compounds) and commercial availability of purified chemicals. Mice received daily topical treatments with compounds at 20% dilutions (w/w) in acetone vehicle for five weeks. Both d,l-limonene and α-pinene had significantly fewer lesions than vehicle application, whereas mice treated with eucalyptol, γ-terpinene, or camphene did not differ from vehicle controls (**Fig. 5-S1**). Thus, we conclude that d,l-limonene and/or α-pinene likely mediate the antitumor effects of CWO.

**Fig. 5.**
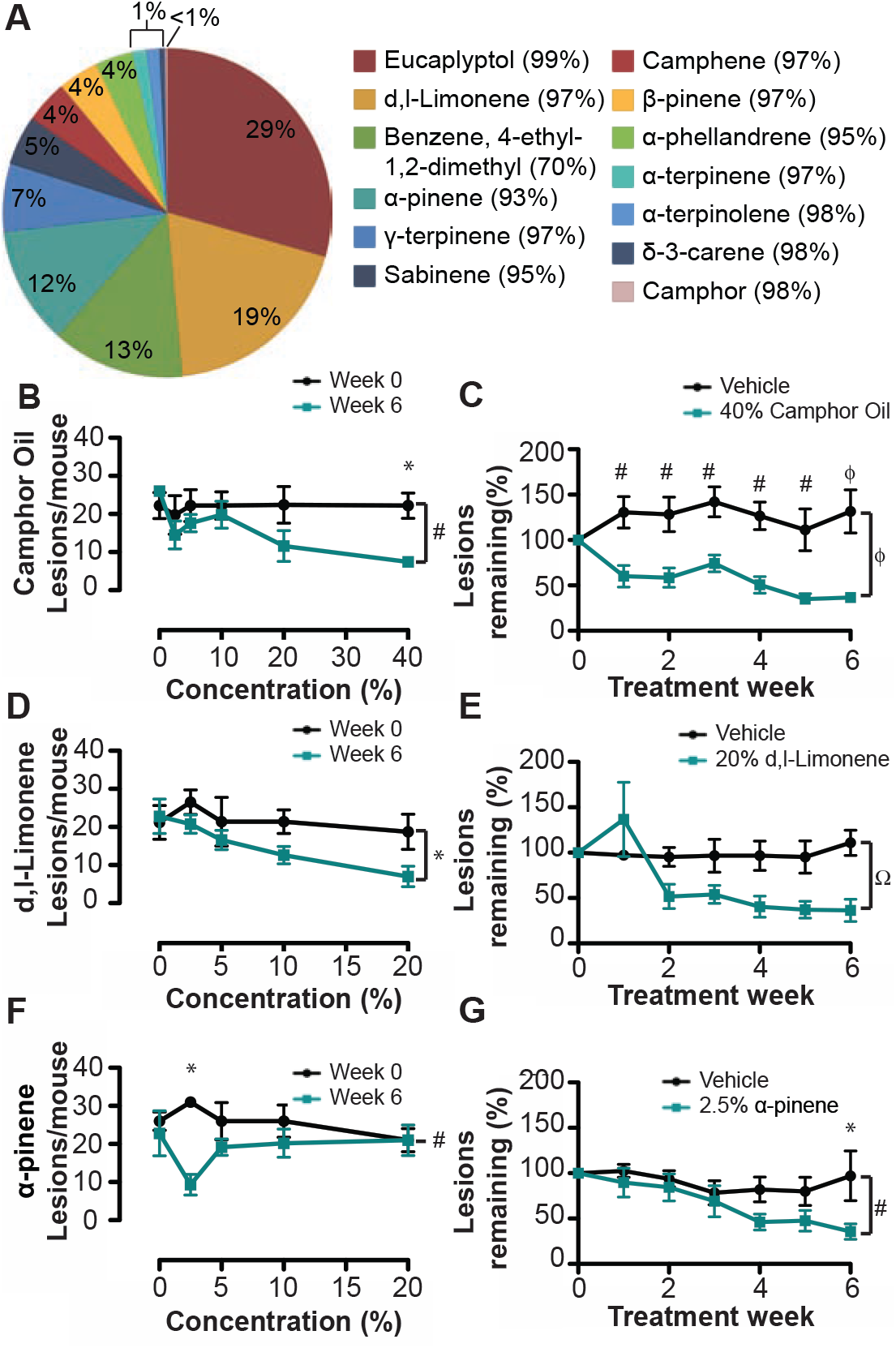
CWO constituents partially recapitulate the effects of CWO. **A.** Gas-chromatography mass spectrometry analysis of CWO. Percentage of constituent present in CWO is shown on the pie chart and the percentage match to reference is shown in parentheses. One compound did not match well to reference. **B-G.** Dose-response studies to identify effective concentrations of CWO and constituents in reducing tumor burden. Tumors were induced as before and mice were split into matched groups (*N*=5 mice/group). Mice were treated for 6 weeks with CWO or constituent terpenes (0–40%). **B,D,F** show the number of lesions/mouse at week 0 (black line) and 6 weeks (teal line) for each concentration tested. **C, E, F** show % of tumors remaining for vehicle (black line) and the optimal concentration of compound (teal line). All data were analyzed with two-way ANOVA with Bonferroni post-hoc. Significance of group P-values is denoted on side of graph, and post-hoc differences at each week are noted above the time point. *P<0.05, ^#^P<0.01, ^Ω^*P*<0.001, ^ϕ^P<0.0001. See **Fig. 5-S1** for pilot experiments with terpenes. **B.** CWO showed a dose dose-dependent effect and 40% CWO was most effective **C.** 40% CWO caused a significant effect after one week of treatment, and was significantly different from vehicle treatment. **D.** d,l-Limonene showed a slight dose-dependent response, with 20% causing the greatest reduction in tumor burden. **E.** 20% d,l-Limonene caused a significant reduction in tumor burden compared to vehicle. **F.** α-Pinene was maximally effective at 2.5% concentration. **G.** 2.5% α-Pinene was significantly different from vehicle after 6 weeks of treatment.

We next sought to identify effective concentrations of terpenes for tumor reduction *in vivo*. Tumor-bearing mice were split into matched groups and treated daily with a range of concentrations of CWO, d,l-limonene, or α-pinene (0–40% N=5 mice per group). CWO was rapidly effective at high concentrations (40%; **Fig. 5B**), with significant reduction in tumor burden achieved within one week (**Fig. 5C**). In pilot studies, 40% solutions of either d,l-limonene or α-pinene produced marked skin irritation or toxicity; thus, these chemicals were applied up to 20% (**Fig. 5D**). D,l-limonene produced the largest reduction in skin tumors at 20%, which was apparent two weeks after the start of treatment (**Fig. 5D-E**). By contrast, 2.5% α-pinene resulted in significant reductions in tumor burden (**Fig. 5F-G**). Interestingly, higher concentrations of α-pinene did not reduce lesions in this cohort. Together, these results indicate that, compared with α-pinene or d,l-limonene, camphor essential oil is better tolerated and leads to more rapid reduction in tumor load. This finding implies that naturally derived CWO contains additional components that work in synergy with d,l-limonene and α-pinene to promote their antitumor effects *in vivo*.

## Discussion

This study identifies a previously unsuspected use for an essential oil to induce regression of keratinocyte-derived malignancies. CWO and its derivatives induced a striking reduction in tumor burden in pre-clinical mouse models *in vivo*. Remarkably, daily topical treatment with CWO stimulated regression of pre-existing lesions and reduced the incidence of malignant conversions by half. Furthermore, CWO induced calcium/calcineurin-dependent NFAT translocation, and this in turn had direct physiological effects on keratinocytes. CWO treatment resulted in a multitude of transcriptional changes that could affect the tumor microenvironment. Specifically, tumor clearance was mediated through cytotoxic CD8+ T cells, arguing for immune-dependent clearance of tumor cells. Collectively, these findings suggest that CWO and its terpene constituents comprise a naturally occurring immune cell modulator.

CWO’s terpene constituents are commonly used in commercial manufacturing, culinary and medicinal applications. For example, eucalyptol is used in mouthwash and cough suppressants. D,l-Limonene is prominent in citrus oils used as flavorings in foods and beverages, and is a solvent in household cleaning products. The scent of pine is conferred by α-pinene in household products, and γ-terpinene is used in cosmetics. Thus, humans are frequently exposed to these compounds, which are well tolerated on skin. We identified α-pinene and d,l-limonene as active compounds that promote tumor regression. Future studies are needed to assess whether these components act synergistically or separately on molecular pathways. The prominence of CWO constituents in common household items poses an interesting avenue for future investigation into whether exposure to these terpenes over a life-span alters the risk for keratinocyte-derived lesions.

A handful of recent studies suggest a role of CWO’s terpene constituents in experimental cancer models (Bayala et al., 2014, Bhattacharjee & Chatterjee, 2013, Chaudhary, Siddiqui, 2012, Chidambara Murthy, Jayaprakasha, 2012, Kusuhara, Urakami, 2012, Lee, Hyun, 2006, Liu et al., 2011, Russin, Hoesly, 1989). Extracts of *C. camphora* and related species are reported to be cytotoxic to human tumor cells *in vitro* (Liu, Chen, 2011, Satyal, Paudel, 2013). Consistent with our observations, D-limonene has been reported to have preventive effects on pre-malignant lesion formation in a mouse model of cSCC (Chaudhary, Siddiqui, 2012). Inhalation of α-pinene has been suggested to reduce melanoma growth in mouse models (Kusuhara, Urakami, 2012). These studies have shown effectiveness of terpenes on the promotion and establishment phases of tumorigenesis. To our knowledge, this study is the first to show that terpenes, including naturally derived CWO, induce regression of existing tumors *in vivo*. Furthermore, the efficacy of treatment is greater when the natural oil is used, as compared with individual terpenes.

Natural products represent a rich source for small molecule therapeutics; however, the molecular targets of these compounds are often elusive (Chidley et al., 2016, Schenone et al.,2013, Shoemaker, 2006, Ziegler et al., 2013). CWO presents challenges for identifying the precise molecular target due to the complexity of the admixture and potentially combinatorial effects of the terpene constituents. CWO stimulated calcium signaling in keratinocytes, followed by NFAT translocation to the nucleus (**Fig. 6**). These are key findings in understanding the mechanism of action for tumor elimination. An obvious question is – what are CWO’s receptors in skin? TRPV3 was our top candidate, as this calcium channel is expressed in keratinocytes and activated by camphor and other terpenes; however, CWO-mediated effects in keratinocytes were comparable in wildtype and TRPV3 knockout mice. Thus, the direct molecular target of CWO remains an open question for future investigation. CWO’s receptors might include other TRP channels or voltage-gated calcium channels (Cui et al., 2017). Alternatively, CWO could stimulate store-operated release through Gq coupled G-protein activation or by store operated calcium entry (e.g. a STIM-Orai dependent pathway) (Cui, Merritt, 2017, Sumit et al., 2015). Our observation that d,l-Limonene and α-pinene have anti-tumor effects provide a starting point for future studies to identify receptors that link CWO’s active ingredients to NFAT signaling.

**Fig. 6.**
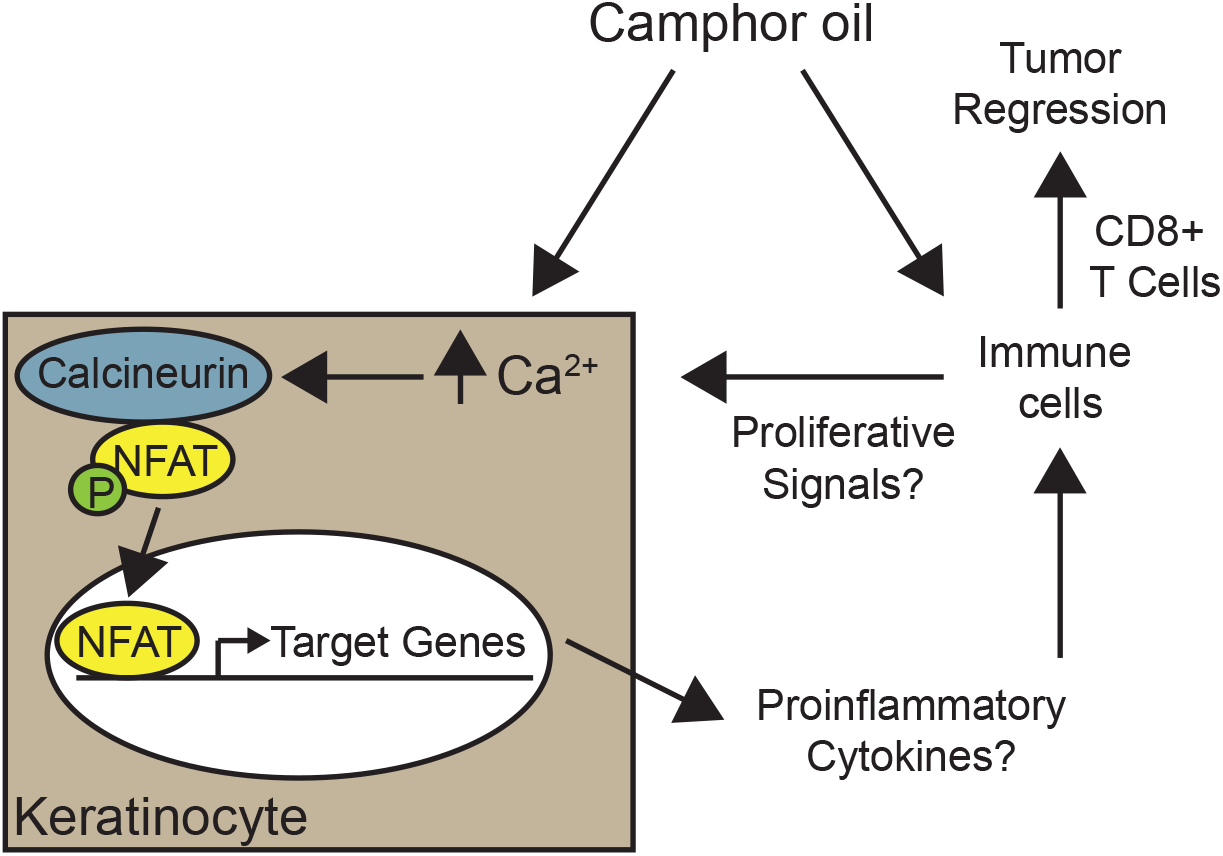
Model for CWO’s mechanism of action. CWO induces calcium/calmodulin dependent NFAT translocation. NFAT induces transcription that in turn alters the tumor microenvironment promoting inflammation and T cell mediated tumor regression.

We propose that CWO stimulates tumor regression in skin through NFAT-dependent signaling and CD8+ T cell-dependent mechanisms (**Fig. 6**). Consistent with the involvement of NFAT signaling, our *in vitro* studies demonstrated that CWO induced calcium signaling and calcium/calcineurin-dependent NFAT translocation in human keratinocytes.

The effects of NFAT signaling in keratinocytes are multifaceted and context dependent. For example, NFAT activation downstream of Notch signaling has been shown to induce a switch from proliferation to differentiation of epidermal keratinocytes (Mammucari, Tommasi di Vignano, 2005, Santini, Talora, 2001). In the hair cycle, NFAT signaling promotes bulge stem cell quiescence by inhibiting the cell cycle regulator CDK4 (Horsley, Aliprantis, 2008, Mammucari, Tommasi di Vignano, 2005, Santini, Talora, 2001). By contrast, overactivation of NFAT in epidermal cells promotes keratinocyte hyperproliferation (Tripathi, Wang, 2014). The latter report is consistent with our finding that CWO increases proliferation of normal human keratinocytes.

How can increased keratinocyte proliferation be reconciled with tumor regression? One possibility is that CWO causes normal keratinocytes to proliferate and outcompete cancer stem cells for metabolic resources. Alternatively, NFAT activation could differentially affect normal and cancer stem cells, promoting proliferation in the former and quiescence in the later (Tripathi, Wang, 2014, Wu, Nguyen, 2010). Collectively, this alteration in proliferative potential could tip the scale to promote tumor regression.

Our finding that immune cells are required for CWO’s antitumor effects *in vivo* favor another model: that CWO promotes NFAT translocation in keratinocytes to induce expression of cytokines that alter the tumor microenvironment, stimulating the clearance of tumor cells through immune cell activation. For example, recent work has identified thymic stromal lymphopoietin (TSLP) release downstream of NFAT activation in keratinocytes (Wilson et al., 2013). TSLP is well known for having a role in promoting allergic inflammation (Ziegler, 2012) and when released from keratinocytes, can promote the atopic march that proceeds to asthma development (Zhang et al., 2009). Interestingly, allergic inflammation can reduce risk of certain cancers while increasing the risk for others (Josephs et al., 2013). Several intriguing studies have shown that TSLP has an additional role in stimulating antitumor immunity through CD4+ and CD8+ T-cell activation (Demehri et al., 2016, Demehri et al., 2012, Di Piazza et al., 2012). TSLP released from keratinocytes can directly stimulate CD4+ T_H_ activation and recruitment of cytotoxic T cells to tumors. Interestingly, we found that TSLP mRNA was upregulated in keratinocytes treated with CWO, indicating this is signaling pathway might contribute to CWO-mediated tumor regression.

Along with effects on keratinocytes, CWO could act directly on T cells, either through resident populations in skin or through systemic effects (Medler & Coussens, 2014, Mueller et al., 2014, Richmond & Harris, 2014). NFAT isoforms have been most heavily studied for their effects on T cells (Hogan, 2017). Upon T cell activation, nuclear NFAT proteins complex with AP1 transcription factors to stimulate gene expression linked to activation. Therefore, CWO could directly influence T cell activation dynamics through NFAT. Future studies will aim to define whether CWO acts directly on keratinocytes, immune cells or both to stimulate anti-tumor immunity.

Collectively, this work identifies CWO as a novel activator of NFAT signaling and calcium/calcineurin-mediated antitumor immunity. Organ transplant recipients that receive long-term treatment with calcineurin inhibitors are at dramatically increased risk for SCC (Dotto, 2011, Goldstein, Fletcher, 2014, Horsley, Aliprantis, 2008, Keyes, Segal, 2013, Yamamoto & Kato, 1994), arguing that NFAT activators might be used to reduce epithelial tumor risk. As these studies investigated the efficacy of CWO and terpene compounds after tumor induction, future studies are needed to determine whether these compounds can also act as a preventive to reduce the development of pre-cancerous lesions. This raises the possibility that CWO could serve as an effective topical treatment to prevent the progression of keratinocyte-derived lesions.

## Materials and Methods

### Animals

Animal use was conducted according to guidelines from the National Institutes of Health's Guide for the Care and Use of Laboratory Animals and the Institutional Animal Care and Use Committee of Columbia University Medical Center. Age-matched female FVB/NJ mice were from Jackson labs were used in all experiments except for TRPV3 knockout mice (Jackson labs stock #010773).

### Tumor induction

Mice were shaved at age 6–7 wks with electric clippers and telogen, the resting hair cycle stage, was confirmed after 2 d. One topical application of 400 nmol DMBA in 200 μL acetone was administered, followed by one week of rest then twice-weekly applications 200 μL of 10 nmol TPA in acetone. This TPA regimen ensured each mouse generated a high tumor burden by 15 wks with at least one malignant conversion before endpoint. Mice were monitored daily and euthanized when they had a tumor >20 mm in diameter, had tumor ulceration leading to loss of skin barrier, showed signs of anemia for ≥24 h, or had a gross appearance indicating distress (hunched posture, lethargy, persistent recumbence).

### Tumor classification criteria

Tumor number and location were documented weekly. A lesion was classified as precancerous based on its appearance as a non-ulcerated, fleshy pedunculated or sessile wart-like mass with a diameter in any dimension ≥2 mm. Lesions were classified as malignant SCCs when they converted to a flattened circular growth with a depressed center, or showed spontaneous ulceration (Allen et al., 2003).

### Treatment paradigm

Mice assigned to the CWO or terpene groups were treated daily with topical 2.5%–40% oil in acetone (wt/wt; 400 μl, applied drop-wise). Control mice were treated with topical acetone vehicle (Veh, 400 μl). Three to seven days after the last TPA treatment, mice were randomly assigned to CWO, terpene, or control treatment groups, matched for total precancerous lesion burden. Mice assigned to the CWO or terpene groups were treated daily with topical 2.5%-40% (wt/wt) oil in acetone (wt/wt; 400 μl applied drop-wise). Mice assigned to the control (acetone vehicle) group were treated daily with topical acetone (400 μl applied drop-wise to lesions). Terpene treatments included CWO (synthetic camphor white oil CAS#8008-51-3 Sigma Aldrich catalog # W223115), Limonene (CAS# 138-86-3 Sigma Aldrich catalog # W524905), α-Pinene (CAS# 80-56-8 Sigma Aldrich catalog # 147524), Eucalyptol (CAS# 470-82-6 Sigma Aldrich Cat# C80601), γ-Terpinene (CAS# 99-85-4 Sigma Aldrich Cat# W355909)

### Cell culture

Primary keratinocytes were derived from discarded foreskin tissue using two-step enzymatic digestion. Connective tissue was removed and skin was treated overnight with dispase at 4 C. Epidermis was separated and treated with 0.25% trypsin and transferred to DMEM with 10% FBS, dissociated with a pipette, filtered with a 70 μm cell strainer and grown in keratinocyte growth media (CnT07, CellNTech). All cells used were <5 passages and plated to 70–85% confluence. At least three independent keratinocyte donors were used in all experiments.

### Calcium imaging

Keratinocytes were loaded with Fura-2 AM ratiometric calcium indicator (5 μM Fura-2 & 1 μM Pluronic in isotonic ringers) for 45 min, washed with isotonic ringers, and allowed to recover for 45 minutes. Data was gathered at 30 frames/min. Baseline recordings of calcium activity were gathered for 2 minutes using MetaFluor software. Cells were treated with 0–0.08% CWO with DMSO vehicle. Cells with baseline calcium ratio greater than two standard deviations above experiment mean were excluded. Mean response greater than 20% of the cell’s baseline calcium ratio were considered responders. Activity was recorded for 5 minutes following compound application, followed by 10 μM histamine as a positive control to ensure that cells were activatable. Analysis was performed using Excel and Prism software. 250–678 cells were analyzed per experiment. Cells were counted as responders if they displayed greater than 20% increase in 340:380 ratio over the cell’s baseline value. The percent of cells that responded to treatment were then calculated per experiment and pooled by treatment condition.

### Cell Viability

Normal human keratinocytes were cultured and plated on an 8-well plate (Lab-Tek II Slide, 8 Chamber (Nalge Nunc International, Cat 154534). Cells were treated with keratinocyte media (live control) and media with CWO (Sigma Aldrich, Cat#W223115, Lot MKBP1241V, 0.01%, 0.02%, 0.04%, 0.08% by volume) for 30 minutes at 37 degrees Celsius. LIVE/DEAD Reduced Biohazard Cell Viability Kit #1, green & red fluorescence (Thermo Fisher Scientific, Cat#L-7013) performed according to manufacturer’s directions to assess cell viability. Cells were imaged on confocal microscope on the same day at 10x. Quantification was performed by counting number of live (green) and dead (red) cells per field. One image was taken per well (N=3 wells per experimental replicate).

### NFAT assays

Keratinocytes were treated with CWO (0-0.04% by volume) for 30 min at 37 C. For CSA treatments, cells were pretreated for 1 h with 1 μM CSA (Fisher 239835) or an equal volume of Veh (ethanol). Cells were fixed with 4% paraformaldehyde in PBS, and stained with NFATc1 (7A6) antibody overnight (Santa Cruz Biotechnology sc-7294), stained with secondary antibody (Alexa Fluor 488, Invitrogen A11029) and mounted with fluoromount + DAPI (Southern Biotech cat 0100–20). Six images per well were taken using a confocal microscope at 40x and translocated/total cells in were quantified in ImageJ. The average fraction of cells/well with nuclear translocation was calculated per experiment.

### Flow cytometry

Mice were shaved and 1.5–2 wks later telogen was confirmed and mice were treated with CSA (Calbiochem, 20 mg/kg, dissolved 10 mg/ml in Veh (10% ethanol, 90% olive oil)) or an equal volume Veh by intraperitoneal injections. A second injection of CSA or Veh was administered 24 h later along with 400 µL topical treatment of CWO or Veh (acetone). Mice were injected with EdU 23 h later (10 uL/g weight of 5 mg/ml EdU mixed in PBS). Mice were sacrificed 1 h later and keratinocytes were isolated according to published protocols (Doucet & Owens, 2015). EdU was detected via the click-it reaction (Invitrogen) and cells were stained with antibodies against cell surface antigens on basal keratinocytes (α6-Integrin, BD-Pharmingen 555736). Cell fluorescence was analyzed on a BD FACSCantoII flow cytometer. The percentage of α6-Integrin+ cells that were positive for EdU in each group were quantified in FlowJo.

### RNA sequencing

10 FVB mice were shaved. One week later, five mice received 400 μL topical 20% CWO. 24 h after treatment, mice were sacrificed and epidermis was isolated via 40-min incubation in 3.8% ammonium thiocyanate (Sigma-Aldrich, CAS 1762-95-4) in RNAse free PBS (Ambion, 10x PBS buffer pH 7.4, PN AM9625). The work area was kept RNAse free using RNaseZap (PN AM9780). The epidermis was homogenized in 1 mL TRizol (Thermo, Cat. #15596026) using Omni International homogenizer with soft tissue disposable tips (PN 32750). RNA isolation proceeded using the TRIzol Plus RNA Purification Kit (Thermo Fisher, Cat. #12183-555). Briefly, homogenized epidermis in TRizol was incubated at room temperature for five minutes. 0.2 mL chloroform (Sigma Aldrich, CAS 67-66-3, PN 360927) was added. Lysate was centrifuged in 5 PRIME, phase lock heavy 2 mL tubes (Cat. # 2303830). An equal volume of ethanol was added (Decon, CAS 64-17-5). DNAse treatment using RNAse-Free DNAse was performed (Qiagen Cat. #79254). RNA was eluted using DEPC Treated Water (Invitrogen, Part no. 46-2224). A series of washing steps was conducted following the directions provided in the TRIzol Plus RNA Purification Kit. The purity of RNA was confirmed using a Bioanalyzer (Instrument Name DE72901373, Firmware C.01.069, Type G2939A). Three samples from each treatment with the best RNA integrity were chosen for RNA sequencing. Sequencing was performed using TruSeq RNA Sample Prep Kit v2 (read length 1×100 bp, read count 30M) at the JP Sulzberger Columbia Genome Center Core. Differential expression between vehicle and CWO groups was performed using DESeq. For epidermal samples, one vehicle control was identified as an outlier and removed from analysis.

RNA-sequencing of tumors was initiated using a published DMBA-TPA protocol as described above. Mice were treated for six weeks and then tumors were harvested. Four pre-malignant tumors of approximately the same size per mouse were pooled (N=3 mice per treatment). The pooled tumors were homogenized in 1 mL TRizol using an Omni International homogenizer with soft tissue disposable tips. RNA extraction proceeded as above.

RNA-Seq data have been deposited in NCBI's Gene Expression Omnibus (Edgar et al., 2002)and are accessible through GEO Series accession number GSE117557 (https://www.ncbi.nlm.nih.gov/geo/query/acc.cgi?acc=GSE117557).

Differential expression was determined using DESeq and data were filtered for genes with mean expression >5 RPKM and p≥0.05. This resulted in 1655 dysregulated genes in skin and 293 dysregulated genes in tumors. GO analysis was performed using Cytoscape (Shannon et al., 2003)with the Clugo plugin (Bindea et al., 2009). Data were analyzed to identify significantly altered GO terms in both datasets using Enrichment/Depletion (two-sided hypergeometric test with a Bonferroni step-down correction). Data shown are GO levels 4–7 with at least 2 genes per cluster representing at least 4% of the genes in the cluster with GO term fusion.

### T-cell blocking

T-cell blocking antibodies were given by intraperitoneal injection 100 μg/injection on treatment day -1, 2, then once weekly until endpoint. Antibodies used were anti-mouse CD8 (Clone 2.43 BE0061), anti-mouse CD4 (Clone GK1.5 BE0003–1), and Rat IgG2b (BE0090) (BioXcell). This treatment paradigm has previously been shown to efficiently knockdown T cells in mice (Fransen et al., 2013).

### Immune cell extraction

Immune cells were isolated from blood to analyze antibody knockdown efficiency. Blood was taken by cardiac puncture and directly put into red blood cell lysis buffer. Blood cells were lysed for 10 minutes, resuspended in DMEM, then stained with antibodies (FITC anti mouse-CD4 GK1.5 eBiosciences 11-0041-82, PE-Cy7 Anti-mouse CD8a eBiosciences 25-0081-82) for 1 h.

### GC-MS analysis

GCMS analysis was performed by NDE Analytical (Pleasonton, CA) using the Agilent GC/MS system 6890/5973 with a TG-624 30 m column length, 1.4 um film thickness, 0.25 mm ID and helium carrier gas. The temperature program used was 100 C for 4 minutes, 100–120 C for 10 minutes, 120–220 C for 6 minutes with a 50 C/minute ramp up rate. Data were matched to existing library spectra.

### Statistics

Data are expressed as mean±SEM unless noted. Statistical analysis was performed using Graphpad (Prism V5). Outliers were identified using Grubbs method with Alpha=1×10^-4^. Graphs showing the number of malignant conversions were fitted using the Boltzmann sigmoidal equation:

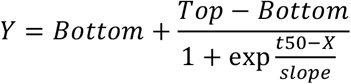

## Author contributions

Conceptualization: YM, AMN, DMO, EAL; Methodology: YM, SAG, EAL, DMO; Validation: YM, LVD; Formal Analysis: YM, SAG, EAL; Investigation: YM, SAG, BAJ, KLM, EAL; Resources: DMO; Data Curation: YM, EAL; Writing-Original Draft: YM; Writing-Review and Editing: BAJ, KLM, LVD, AMN, DMO, EAL; Visualization: YM, SAG, EAL; Supervision: EAL; Project Administration: EAL; Funding Acquisition: DMO, EAL

## Acknowledgements

We thank Yan Lu and Milda Stanislaukas for assistance with histology, Rong Du for assistance with cell culture, Spandan Shah and Siu-Hong Ho for flow cytometry, Dr. Diana Baustista and Caroline Walsh for helpful discussions, Dr. Masashi Natakani for preliminary studies of TRPV3 knockout mice, and Lan Li for critical reading of the manuscript. Funding was provided by NIH/NIAMS R01AR051219 (to EAL) and a pilot award from NIH/NIEHS (P30ES009089). Cell culture and microscopy were performed with support from the Columbia University Skin Disease Resource-Based Center (epiCURE, P30AR069632). RNA sequencing was performed with support from the JP Sulzberger Columbia Genome Center Core (P30CA013696). Flow cytometry was performed using the Columbia Center for Translational Immunology (CCTI) Core facility (P30CA013696). SAG was funded by NIH/NIAMS P30AR044535-14, Columbia University Dean’s Research Fellowship and NIH/NHLBI 5T35HL007616. YM was funded by NIH/NHBLI 1T32HL120826.

## Figure supplements

**Fig. 1-S1.**
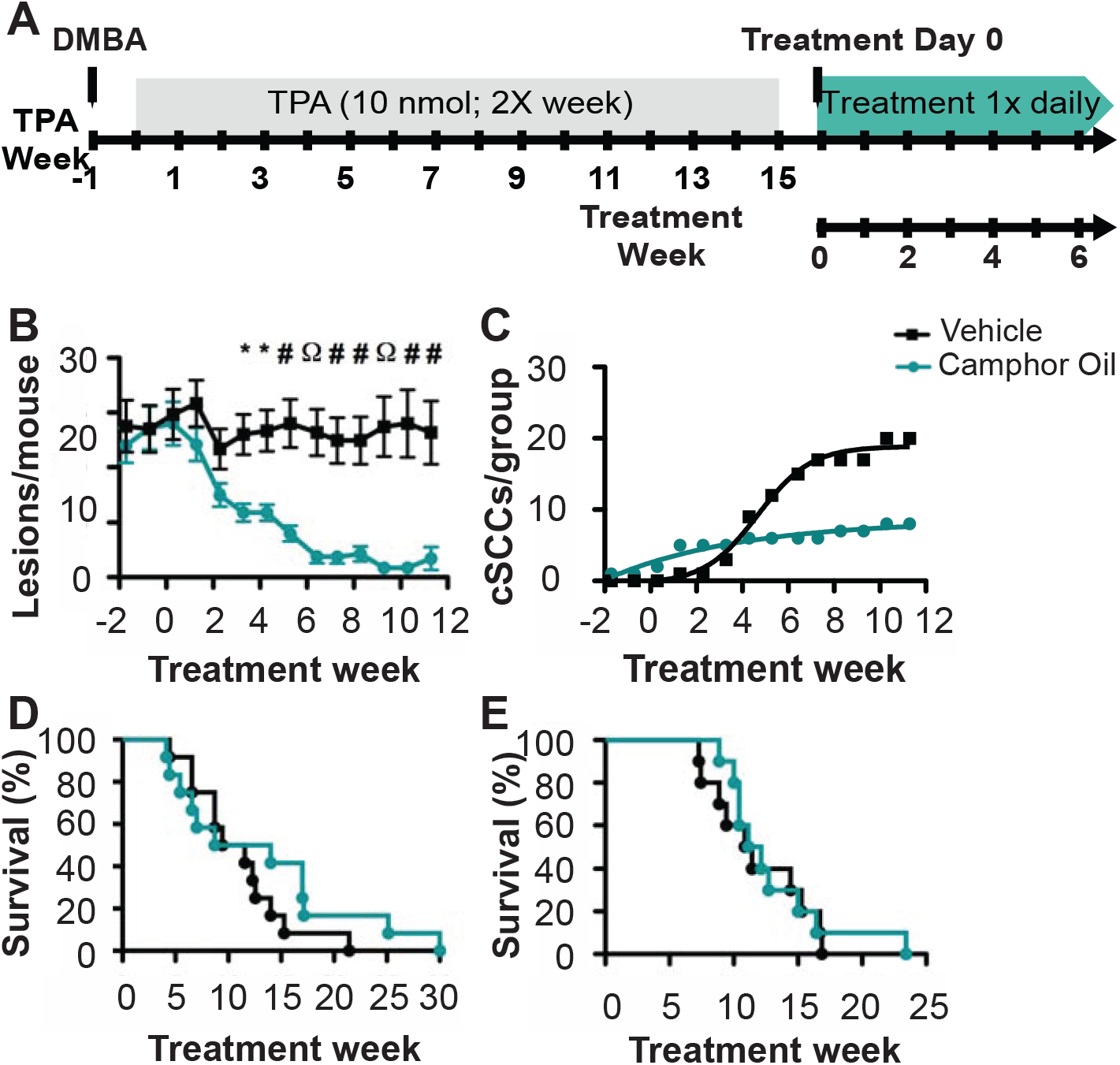
CWO reduces tumor burden in an independent cohort. **A.** Mice were treated once with DMBA to induce mutations followed by twice weekly TPA (10 nM) for 15 weeks. Animals were then split into matched cohorts and treated daily with topical CWO (20%) in vehicle (acetone) or vehicle only. **B.** Topical CWO treatment significantly reduces the total lesion burden (CWO=teal, vehicle=black). Two-way ANOVA [N=12 mice per group, P<0.0001 F(1,252)=106.11]. *P<0.05, ^#^P<0.01, ^Ω^P<0.001 Bonferroni post hoc. **C.** Fewer malignant SCCs formed with CWO treatment than with vehicle treatment. Boltzmann fits, difference between fits F(4,20)=122.4 P<0.0001; CWO: R^2^=0.93, Max cSCC=8.5 weeks; Vehicle: R^2^=0.99, Max cSCC=18.89. **C.** Kaplan-Meyer Survival analysis of CWO and vehicle treated groups reveals no significant difference in survival curves (median CWO=11.36, vehicle=10.5, Log-rank P=0.29), but longer overall survival time for a subset of treated mice. **D.** Similar survival results were found in the first cohort (Fig 1). Median CWO= 11.64, vehicle=11.15, Log-rank P=0.74).

**Fig. 1-S2.**
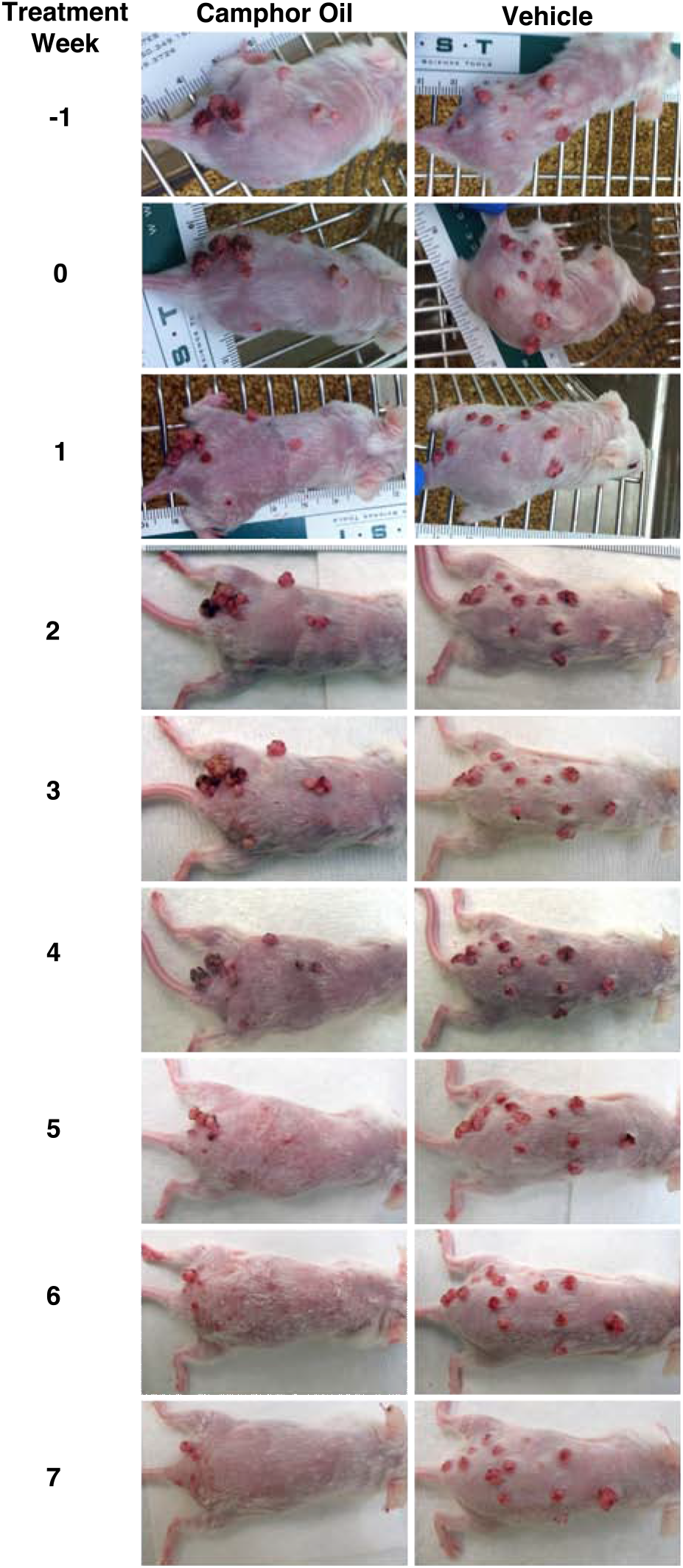
CWO promotes reduction in existing skin lesions through progressive tumor loss. Examples images of single CWO (left) and vehicle (right) treated animals are shown over seven weeks of treatment in a second independent cohort. With CWO treatment, tumors progressively regressed (Week 0 = 10 tumors, Week 7 = 0 tumors); whereas in control mice, tumors were stable in number and grew in size and grade (Week 0 = 13 tumors, Week 7 = 13 tumors).

**Fig. 1-S3.**
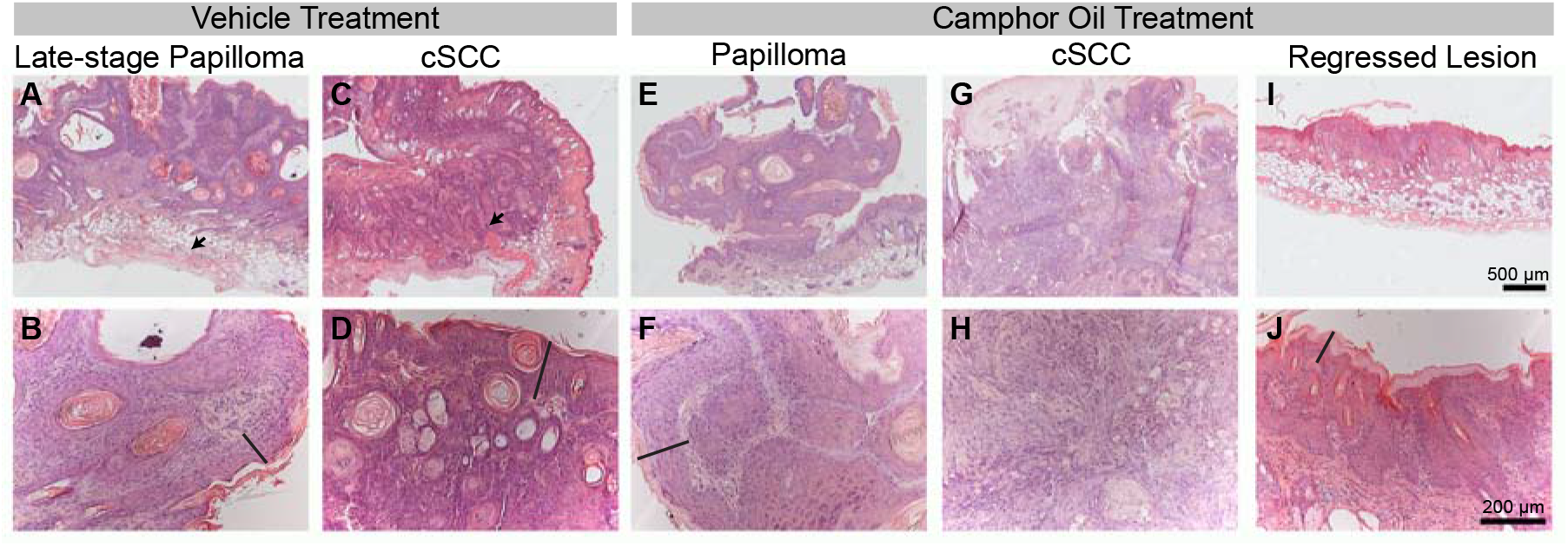
CWO and vehicle tumors appear similar by histology. **A, B.** A late-stage papilloma from a vehicle treated mouse is shown. Note the largely intact layering of the epidermal-dermal boundary (B, black line), panniculus carnosus (arrow) and dermal fat. **C,D.** A fully-converted cSCC from a vehicle-treated animal. Notice the loss of epidermal layering (D, line), and the loss of integrity of the panniculus carnosus (C, arrow). **E, F.** A papilloma from a CWO treated mouse is shown with intact layering of the epidermis (F). **G,H**. A cSCC from a CWO treated mouse is shown. Note the lack of organization in H. **I,J.** Histology from a regressed lesion in a CWO treated mouse is shown. Here, the epidermal layer is thick (line), a sign of hyperproliferation, but normal skin layering seems to be intact.

**Fig. 2-S1.**
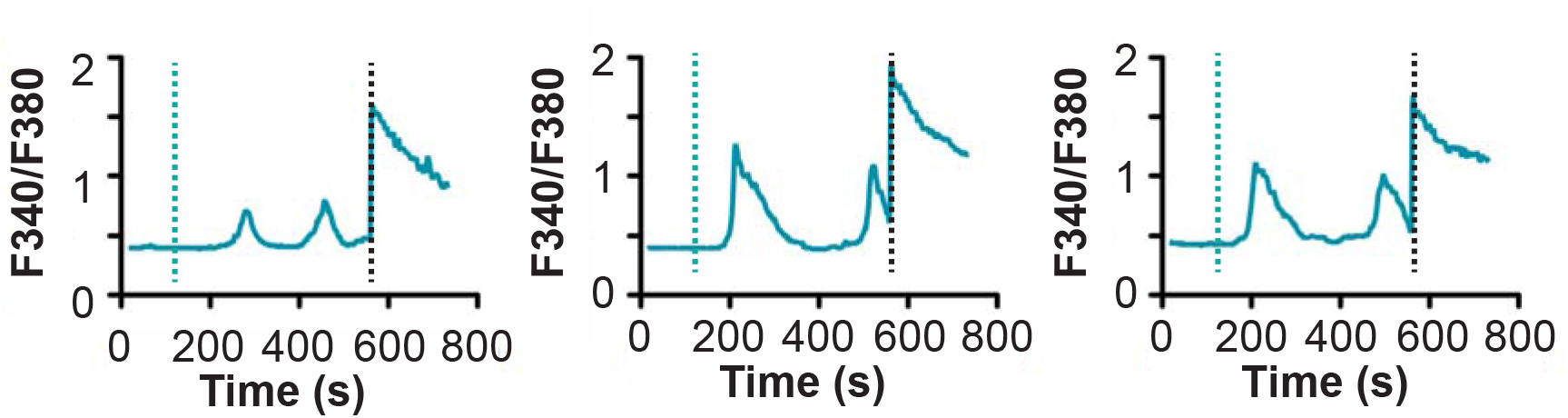
CWO induces waves of calcium signaling in human keratinocytes. Representative example traces show recurrent calcium waves after 0.04% CWO treatment. Teal dashed line indicates CWO application. Black dashed line indicates histamine control.

**Fig. 2-S2.**
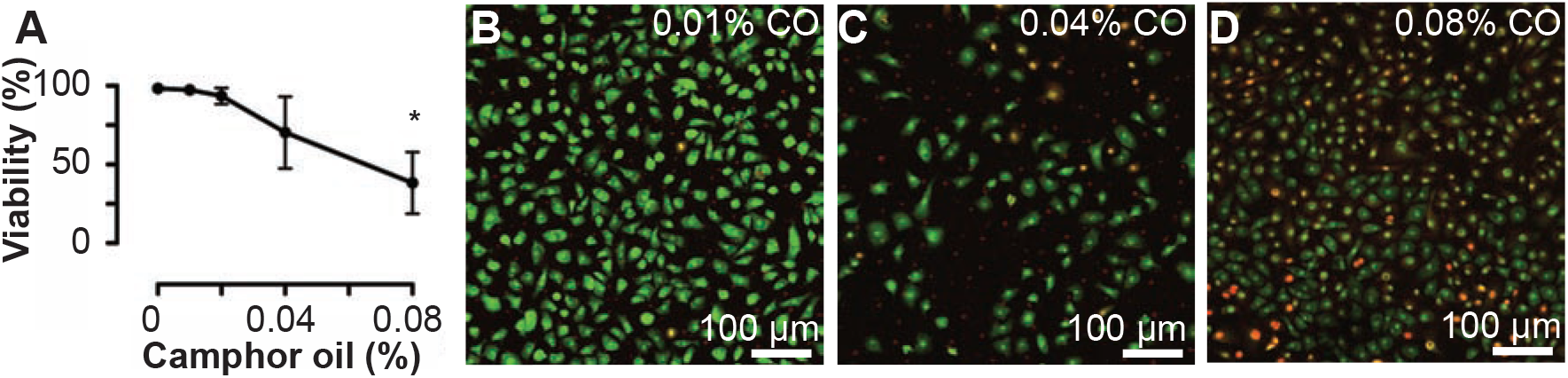
Cultured keratinocytes are viable with CWO treatment. **A.** Percent of normal human epidermal keratinocytes that are viable following 30-minute incubation with 0–0.08% CWO in keratinocyte media (N=3 experiments with >3800 cells per replicate; One-way ANOVA p<0.05, Dunnet’s Multiple Comparison Test against 0% *P<0.05). **B-C**. Representative images from viability assay showing calcein-AM (green) and ethidium homodimer-1 (red) of normal human epidermal keratinocytes treated with 0.01% CWO (**B**) 0.04% CWO (**C**) and 0.08% CWO. All cells are labeled by calcein, but only dead cells by ethidium homodimer-1.

**Fig. 4-S1.**
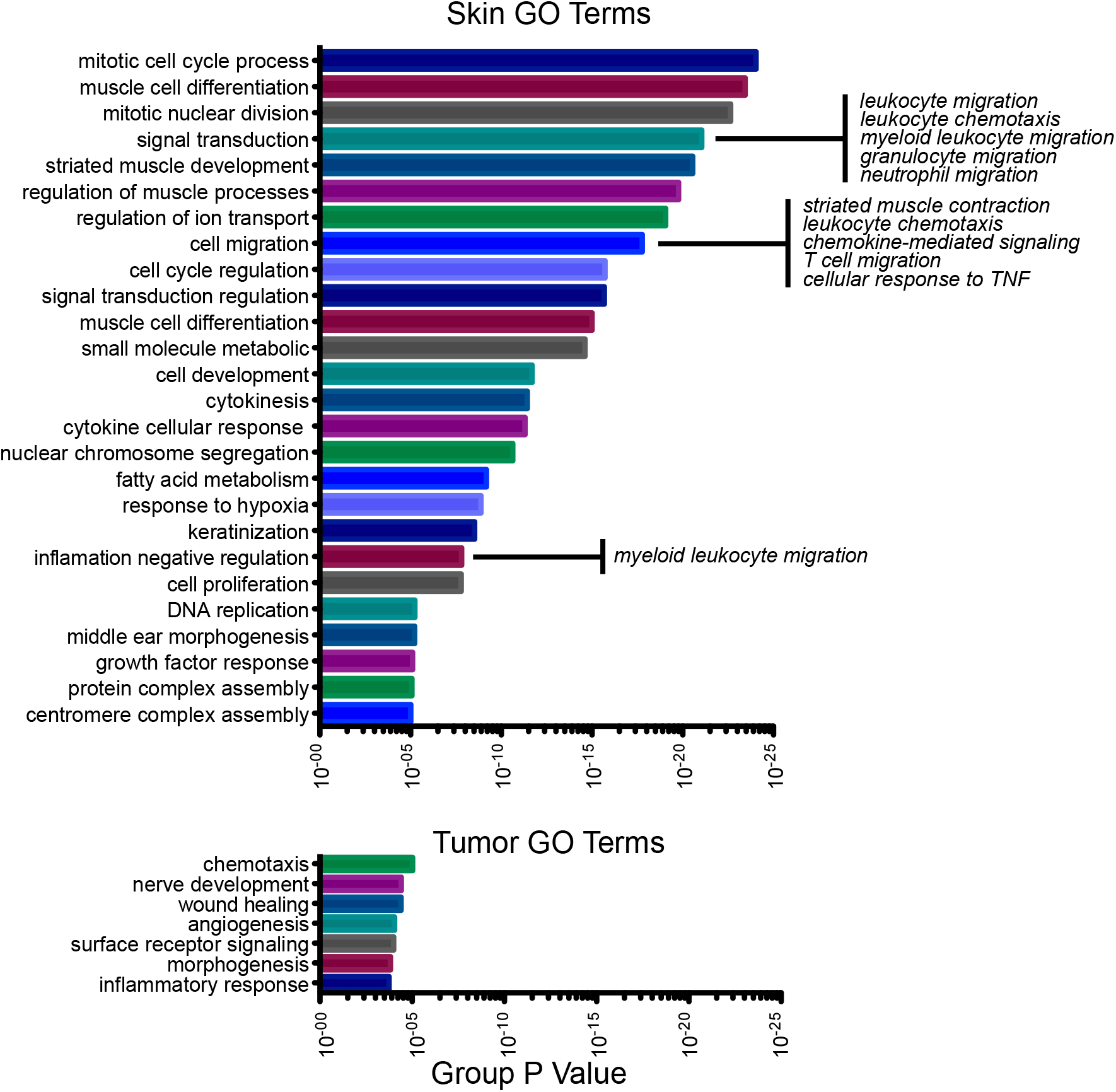
Extended RNA-Seq analysis of tumors and skin treated with CWO. GO analysis of genes differentially expressed in epidermis and tumors treated with CWO compared to vehicle. GO term fusion was used, and selected GO terms within a fused node related to inflammation are shown.

**Fig. 4-S2.**
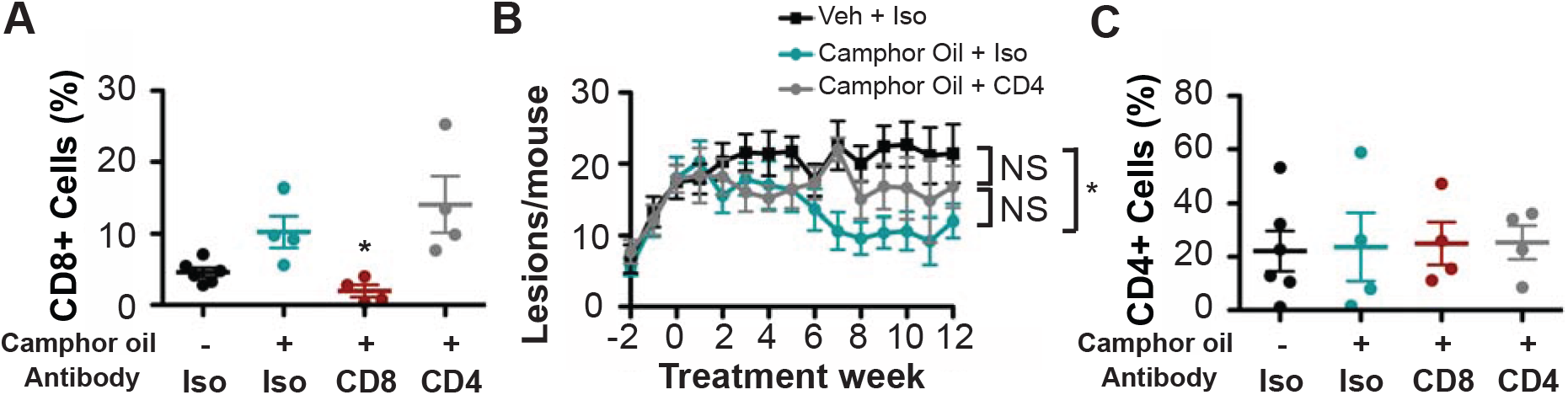
Antibody depletion. Blood from animals in T cell blocking experiments were analyzed for antibody efficiency at sacrifice. Mice were sacrificed between 12–13 weeks after the start of treatment, and 5–6 days after the final antibody injection. **A**. A significant effect of treatment was found on circulating CD8+ T cells (P<0.01) Circulating CD8+ cells were significantly reduced with CD8 antibody treatments. Interestingly, circulating CD8+ cells were increased with CWO treatment. **B**. Co-treatment with CD4 blocking antibodies and topical CWO resulted in an intermediate effect on tumor reduction. CD4/CWO treatment was not significantly (NS) different from either Isotype/CWO or Isotype/veh groups by two-way ANOVA. CWO treatment had similar effects on tumor burden to previous cohorts, *P<0.05 **C**. No effect of treatment was found on circulating CD4+ T cells, indicating that CD4+ T cells were not sufficiently blocked between injections. N=4–6 mice per group.

**Fig. 5-S1.**
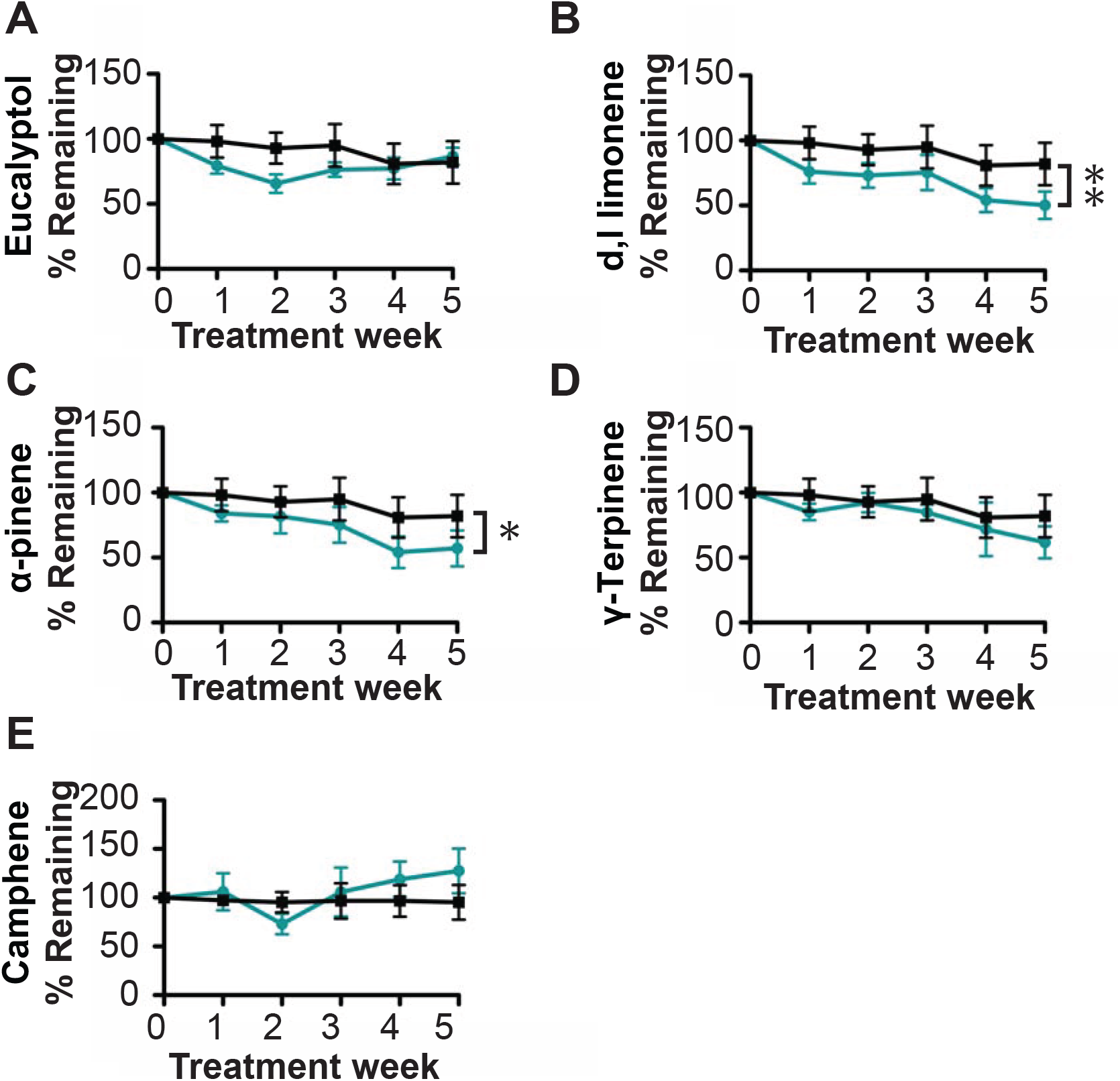
CWO constituents reduce tumor burden. Pilot studies were performed to identify CWO constituents that reduce tumor burden. Tumors were induced as before and mice were split into groups with equal total tumor burdens. Mice were treated with 20% Eucalyptol (**A**), d,l-Limonene (**B**), α-pinene (**C**), g-terpinene (**D**), and camphene (**E**). Data are baseline corrected to tumor counts at time zero for each mouse. Two-way ANOVAs reveal significant treatment effects of d,l-limonene (P<0.01) and α-pinene (P<0.05).

## References

Abel EL, Angel JM, Kiguchi K, DiGiovanni J.Multi-stage chemical carcinogenesis in mouse skin: fundamentals and applications. Nat Protoc. 2009;4:1350–62.

Allen SM, Florell SR, Hanks AN, Alexander A,Diedrich MJ, Altieri DC, et al. Survivin expression in mouse skin prevents papilloma regression and promotes chemical-induced tumor progression. Cancer Res. 2003;63:567–72.

Bachelor MA, Lu Y, Owens DM. L-3-Phosphoserine phosphatase (PSPH) regulates cutaneous squamous cell carcinoma proliferation independent of L-serine biosynthesis. J Dermatol Sci. 2011;63:164–72.

Bakkali F, Averbeck S, Averbeck D, Idaomar M. Biological effects of essential oils--a review. Food Chem Toxicol. 2008;46:446–75.

Bayala B, Bassole IH, Gnoula C, Nebie R, Yonli A, Morel L, et al. Chemical composition, antioxidant, anti-inflammatory and anti-proliferative activities of essential oils of plants from burkina faso. PLoS One. 2014;9:e92122.

Bhattacharjee B, Chatterjee J. Identification of proapoptopic, anti-inflammatory, antiproliferative, anti-invasive and anti-angiogenic targets of essential oils in cardamom by dual reverse virtual screening and binding pose analysis. Asian Pac J Cancer Prev. 2013;14:3735–42.

Bindea G, Mlecnik B, Hackl H, Charoentong P, Tosolini M, Kirilovsky A, et al. ClueGO: a Cytoscape plug-in to decipher functionally grouped gene ontology and pathway annotation networks. Bioinformatics. 2009;25:1091–3.

Blanpain C, Fuchs E. Epidermal stem cells of the skin. Annu Rev Cell Dev Biol. 2006;22:339–73.

Borbiro I, Lisztes E, Toth BI, Czifra G, Olah A, Szollosi AG, et al. Activation of transient receptor potential vanilloid-3 inhibits human hair growth. J Invest Dermatol. 2011;131:1605–14.

Chaudhary SC, Siddiqui MS, Athar M, Alam MS.D-Limonene modulates inflammation, oxidative stress and Ras-ERK pathway to inhibit murine skin tumorigenesis. Hum Exp Toxicol. 2012;31:798–811.

Chidambara Murthy KN, Jayaprakasha GK, Patil BS. D-limonene rich volatile oil from blood oranges inhibits angiogenesis, metastasis and cell death in human colon cancer cells. Life Sci. 2012;91:429–39.

Chidley C, Trauger SA, Birsoy K, O'Shea EK. The anticancer natural product ophiobolin A induces cytotoxicity by covalent modification of phosphatidylethanolamine. Elife. 2016;5.

Crabtree GR, Olson EN. NFAT signaling: choreographing the social lives of cells. Cell. 2002;109 Suppl:S67–79.

Cui C, Merritt R, Fu L, Pan Z. Targeting calcium signaling in cancer therapy. Acta Pharm Sin B. 2017;7:3–17.

Demehri S, Cunningham TJ, Manivasagam S, Ngo KH, Moradi Tuchayi S, Reddy R, et al. Thymic stromal lymphopoietin blocks early stages of breast carcinogenesis. J Clin Invest. 2016;126:1458–70.

Demehri S, Turkoz A, Manivasagam S, Yockey LJ, Turkoz M, Kopan R. Elevated epidermal thymic stromal lymphopoietin levels establish an antitumor environment in the skin. Cancer Cell. 2012;22:494–505.

Di Piazza M, Nowell CS, Koch U, Durham AD, Radtke F. Loss of cutaneous TSLP-dependent immune responses skews the balance of inflammation from tumor protective to tumor promoting. Cancer Cell. 2012;22:479–93.

Dodds A, Chia A, Shumack S. Actinic keratosis: rationale and management. Dermatol Ther (Heidelb). 2014;4:11–31.

Dotto GP. Calcineurin signaling as a negative determinant of keratinocyte cancer stem cell potential and carcinogenesis Cancer Res. 2011;71:2029–33.

Doucet YS, Owens DM. Isolation and functional assessment of cutaneous stem cells. Methods Mol Biol. 2015;1235:147–64.

Edgar R, Domrachev M, Lash AE. Gene Expression Omnibus: NCBI gene expression and hybridization array data repository. Nucleic Acids Res. 2002;30:207–10.

Euvrard S, Kanitakis J, Claudy A. Skin cancers after organ transplantation. N Engl J Med. 2003;348:1681–91.

Fransen MF, van der Sluis TC, Ossendorp F, Arens R, Melief CJ. Controlled local delivery of CTLA-4 blocking antibody induces CD8+ T-cell-dependent tumor eradication and decreases risk of toxic side effects. Clin Cancer Res. 2013;19:5381–9.

Goldstein J, Fletcher S, Roth E, Wu C, Chun A, Horsley V. Calcineurin/Nfatc1 signaling links skin stem cell quiescence to hormonal signaling during pregnancy and lactation. Genes Dev. 2014;28:983–94.

Hannanta-Anan P, Chow BY. Optogenetic Control of Calcium Oscillation Waveform Defines NFAT as an Integrator of Calcium Load. Cell Syst. 2016;2:283–8.

Hennings H, Glick AB, Lowry DT, Krsmanovic LS, Sly LM, Yuspa SH. FVB/N mice: an inbred strain sensitive to the chemical induction of squamous cell carcinomas in the skin. Carcinogenesis. 1993;14:2353–8.

Hogan PG. Calcium-NFAT transcriptional signalling in T cell activation and T cell exhaustion. Cell Calcium. 2017.

Horsley V, Aliprantis AO, Polak L, Glimcher LH, Fuchs E. NFATc1 balances quiescence and proliferation of skin stem cells. Cell. 2008;132:299–310.

Josephs DH, Spicer JF, Corrigan CJ, Gould HJ, Karagiannis SN. Epidemiological associations of allergy, IgE and cancer. Clin Exp Allergy. 2013;43:1110–23.

Keyes BE, Segal JP, Heller E, Lien WH, Chang CY, Guo X, et al. Nfatc1 orchestrates aging in hair follicle stem cells. Proc Natl Acad Sci U S A. 2013;110:E4950–9.

Kusuhara M, Urakami K, Masuda Y, Zangiacomi V, Ishii H, Tai S, et al. Fragrant environment with alpha-pinene decreases tumor growth in mice. Biomed Res. 2012;33:57–61.

Lee HJ, Hyun EA, Yoon WJ, Kim BH, Rhee MH, Kang HK, et al. In vitro anti-inflammatory and anti-oxidative effects of Cinnamomum camphora extracts. J Ethnopharmacol. 2006;103:208–16.

Leonardi-Bee J, Ellison T, Bath-Hextall F. Smoking and the risk of nonmelanoma skin cancer: systematic review and meta-analysis. Arch Dermatol. 2012;148:939–46.

Li W, Llopis J, Whitney M, Zlokarnik G, Tsien RY. Cell-permeant caged InsP3 ester shows that Ca2+ spike frequency can optimize gene expression. Nature. 1998;392:936–41.

Liu CH, Chen CY, Huang AM, Li JH. Subamolide A, a component isolated from Cinnamomum subavenium, induces apoptosis mediated by mitochondria-dependent, p53 and ERK1/2 pathways in human urothelial carcinoma cell line NTUB1. J Ethnopharmacol. 2011;137:503–11.

Mammucari C, Tommasi di Vignano A, Sharov AA, Neilson J, Havrda MC, Roop DR, et al. Integration of Notch 1 and calcineurin/NFAT signaling pathways in keratinocyte growth and differentiation control. Dev Cell. 2005;8:665–76.

Medler TR, Coussens LM. Duality of the immune response in cancer: lessons learned from skin. J Invest Dermatol. 2014;134:E23–8.

Mueller SN, Zaid A, Carbone FR. Tissue-resident T cells: dynamic players in skin immunity. Front Immunol. 2014;5:332.

Nassar D, Latil M, Boeckx B, Lambrechts D, Blanpain C. Genomic landscape of carcinogen-induced and genetically induced mouse skin squamous cell carcinoma. Nat Med. 2015;21:946–54.

Newman DJ, Cragg GM. Natural Products as Sources of New Drugs from 1981 to 2014. J Nat Prod. 2016;79:629–61.

Owens DM, Wei S, Smart RC. A multihit, multistage model of chemical carcinogenesis. Carcinogenesis. 1999;20:1837–44.

Oz M, Lozon Y, Sultan A, Yang KH, Galadari S. Effects of monoterpenes on ion channels of excitable cells. Pharmacol Ther. 2015;152:83–97.

Richmond JM, Harris JE. Immunology and skin in health and disease. Cold Spring Harb Perspect Med. 2014;4:a015339.

Rodrigues T, Sieglitz F, Bernardes GJ. Natural product modulators of transient receptor potential (TRP) channels as potential anti-cancer agents. Chem Soc Rev. 2016;45:6130–7.

Russin WA, Hoesly JD, Elson CE, Tanner MA, Gould MN. Inhibition of rat mammary carcinogenesis by monoterpenoids. Carcinogenesis. 1989;10:2161–4.

Santini MP, Talora C, Seki T, Bolgan L, Dotto GP. Cross talk among calcineurin, Sp1/Sp3, and NFAT in control of p21(WAF1/CIP1) expression in keratinocyte differentiation. Proc Natl Acad Sci U S A. 2001;98:9575–80.

Satyal P, Paudel P, Poudel A, Dosoky NS, Pokharel KK, Setzer WN. Bioactivities and compositional analyses of Cinnamomum essential oils from Nepal: C. camphora, C. tamala, and C. glaucescens. Nat Prod Commun. 2013;8:1777–84.

Schenone M, Dancik V, Wagner BK, Clemons PA. Target identification and mechanism of action in chemical biology and drug discovery. Nat Chem Biol. 2013;9:232–40.

Shannon P, Markiel A, Ozier O, Baliga NS, Wang JT, Ramage D, et al. Cytoscape: a software environment for integrated models of biomolecular interaction networks. Genome Res. 2003;13:2498–504.

Shiku H. Importance of CD4+ helper T-cells in antitumor immunity. Int J Hematol. 2003;77:435–8.

Shoemaker RH. The NCI60 human tumour cell line anticancer drug screen. Nat Rev Cancer. 2006;6:813–23.

Sumit M, Neubig RR, Takayama S, Linderman JJ. Band-pass processing in a GPCR signaling pathway selects for NFAT transcription factor activation. Integr Biol (Camb). 2015;7:1378–86.

Tripathi P, Wang Y, Coussens M, Manda KR, Casey AM, Lin C, et al. Activation of NFAT signaling establishes a tumorigenic microenvironment through cell autonomous and non-cell autonomous mechanisms. Oncogene. 2014;33:1840–9.

Vogt-Eisele AK, Weber K, Sherkheli MA, Vielhaber G, Panten J, Gisselmann G, et al. Monoterpenoid agonists of TRPV3. Br J Pharmacol. 2007;151:530–40.

Wilson SR, The L, Batia LM, Beattie K, Katibah GE, McClain SP, et al. The epithelial cell-derived atopic dermatitis cytokine TSLP activates neurons to induce itch. Cell. 2013;155:285–95.

Wu X, Nguyen BC, Dziunycz P, Chang S, Brooks Y, Lefort K, et al. Opposing roles for calcineurin and ATF3 in squamous skin cancer. Nature. 2010;465:368–72.

Yamamoto S, Kato R. Hair growth-stimulating effects of cyclosporin A and FK506, potent immunosuppressants. J Dermatol Sci. 1994;7 Suppl:S47–54.

Yang F, Long E, Wen J, Cao L, Zhu C, Hu H, et al. Linalool, derived from Cinnamomum camphora (L.) Presl leaf extracts, possesses molluscicidal activity against Oncomelania hupensis and inhibits infection of Schistosoma japonicum. Parasit Vectors. 2014;7:407.

Yu SH, Bordeaux JS, Baron ED. The immune system and skin cancer. Adv Exp Med Biol. 2014;810:182–91.

Zhang Z, Hener P, Frossard N, Kato S, Metzger D, Li M, et al. Thymic stromal lymphopoietin overproduced by keratinocytes in mouse skin aggravates experimental asthma. Proc Natl Acad Sci U S A. 2009;106:1536–41.

Ziegler S, Pries V, Hedberg C, Waldmann H. Target identification for small bioactive molecules: finding the needle in the haystack. Angew Chem Int Ed Engl. 2013;52:2744–92.

Ziegler SF. Thymic stromal lymphopoietin and allergic disease. J Allergy Clin Immunol. 2012;130:845–52.

